# Screening of Botanical Drugs Against Lassa Virus Entry

**DOI:** 10.1101/2020.07.24.220749

**Authors:** Yang Liu, Jiao Guo, Junyuan Cao, Guangshun Zhang, Xiaoying Jia, Peilin Wang, Gengfu Xiao, Wei Wang

## Abstract

Lassa virus (LASV) belongs to the Old World *Mammarenavirus* genus (family *Arenaviridae*) and is classified as a category A biological threat agent. At present, there are no approved drugs or vaccines specific for LASV. In this study, high-throughput screening of a botanical drug library was performed against LASV entry using a pseudotype virus bearing the LASV envelope glycoprotein (GPC). Two hit compounds, bergamottin and casticin, were identified as LASV entry inhibitors in the micromolar range. A mechanistic study revealed that casticin inhibited LASV entry by blocking low pH-induced membrane fusion. Adaptive mutant analyses demonstrated that the F446L mutation, located in the transmembrane domain of GP2, conferred resistance to casticin. Furthermore, casticin extended its antiviral spectrum to the New World (NW) pathogenic mammarenaviruses, and mutation of the conserved F446 conferred NW resistance to casticin. Unlike casticin, bergamottin has little effect on LASV GPC-mediated membrane fusion, while it inhibited LASV entry by blocking endocytic trafficking. Our study shows that both bergamottin and casticin are candidates for LASV therapy, indicating that the conserved F446 plays important roles in drug resistance in mammarenaviruses.

**IMPORTANCE:** Currently, there is no approved therapy to treat Lassa fever (LASF); we aimed to find candidates for LASF therapy. Herein, we screened a botanical drug library and identified two compounds, bergamottin and casticin, that inhibited LASV entry via different mechanisms.

## INTRODUCTION

Lassa virus (LASV) is an enveloped, negative-sense, bisegmented RNA virus belonging to the *Mammarenavirus* genus (family *Arenaviridae*). Mammarenaviruses include 39 unique species currently recognized by the International Committee on Taxonomy of Viruses (1). The original classification of mammarenaviruses, based mainly on virus genetics, serology, antigenic properties, and geographical relationships, divided them into New World (NW) and Old World (OW) mammarenaviruses (2). The OW LASV and Lujo virus (LUJV), as well as some NW mammarenaviruses, including the Junín virus (JUNV), Machupo virus (MACV), Guanarito virus (GTOV), Chapare virus (CHAPV), and Sabiá virus (SBAV), are known to cause severe hemorrhagic fever and are listed as biosafety level (BSL) 4 agents (3, 4).

Mammarenavirus RNA genomes encode the viral polymerase, nucleoprotein, matrix protein (Z), and glycoprotein complex (GPC). GPC is synthesized as a polypeptide precursor that is sequentially cleaved by a signal peptidase and the cellular protease subtilisin kexin isozyme-1/site-1 protease to generate the three subunits of the mature complex: the retained stable-signal peptide (SSP), the receptor-binding subunit GP1, and the membrane fusion subunit GP2 (2, 5-7). The three non-covalently bound subunits SSP, GP1 and GP2 form a (SSP/GP1/GP2)_3_ trimeric complex that presents at the surface of the mature virion and plays essential roles in virus entry. The unusual retained SSP interacts with the membrane proximal external region as well as the transmembrane domain of GP2, and thus stabilizes the prefusion conformation of GPC and provides an interface targeted by entry inhibitors (8-10).

Lassa hemorrhagic fever is epidemic annually in West Africa, and peaks during the dry seasons. In 2020, Nigeria faced the largest outbreak with 4,761 suspected cases, and 1,006 confirmed as of May 17, according to the Nigeria Centre for Disease Control. To date, no vaccines or specific antiviral agents against LASV are available. Therapeutic strategies are limited to the administration of ribavirin in the early course of the illness. To address this issue, we screened a botanical drug library of 1,034 compounds. All the compounds were extracted from botanicals with a purity of ≥ 98%. By using a pseudotype of LASV within biosafety level-2 (BSL-2) facilities, we identified that the compounds bergamottin and casticin inhibit LASV entry.

## RESULTS

### Bergamottin and casticin blocked LASV entry

To construct viral systems affording studies of viral entry in BSL-2 laboratories and facilitate high through-output screening, recombinant LASV (LASVrv) based on the vesicular stomatitis virus (VSV) backbone containing the GPF, as well as the LASV GPC gene, were constructed. LASVrv was replication competent at a titer of 1.6×10^7^ PFU/ml. Pseudotype of LASV (LASVpv) was also constructed with the VSV backbone, and as the GPC gene was substituted with the luciferase gene, LASVpv was competent for a single round of viral entry and infection (9).

As shown in Fig. 1A, bergamottin and casticin were selected as hit compounds after three rounds of screening as both compounds exhibited robust inhibition on the LASVrv whereas they had mild effects on VSV, suggesting that they inhibit LASV infection on the entry step. The IC_50_ values of bergamottin and casticin against LASVpv on Vero cells were 3.615 and 0.6954 µM, respectively, while the values obtained for A549 cells were 4.300 and 1.696 µM, respectively, indicating that both compounds exhibit similar inhibition effects on different cell types (Fig 1C and 1E).

**Fig. 1.**
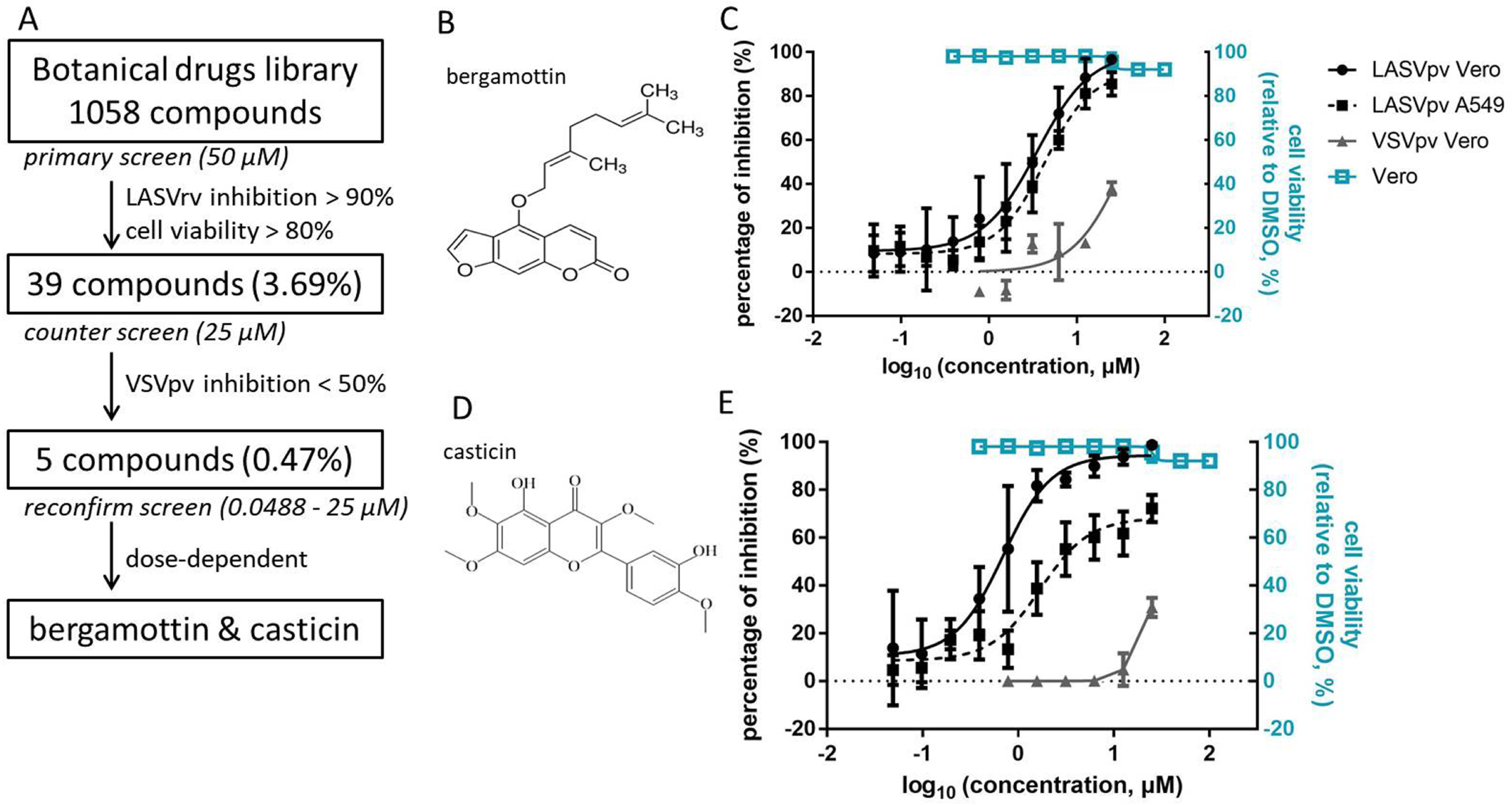
High-throughput screening (HTS) for inhibitors of Lassa virus (LASV) entry from a botanical drug library. (A) HTS assay flowchart. (B) Structure of bergamottin. (C) Dose-response curves of bergamottin. (D) Structure of casticin. (E) Dose-response curves of casticin. Data are represented as the means ± standard deviations (SDs) from six independent experiments.

### Casticin inhibited LASV GPC-mediated membrane fusion

As the unique SSP-GP2 interface played critical roles in stabilizing the pre-fusion conformation of GPC, and thus provided the “Achilles heel” targeted by most mammarenavirus entry inhibitors, the effects of bergamottin and casticin on LASV GPC-mediated membrane fusion were studied. As shown in Fig. 2A, casticin could effectively block low-pH triggered membrane fusion in a dose-dependent manner, while bergamottin showed little effect at all tested concentrations.

**Fig. 2.**
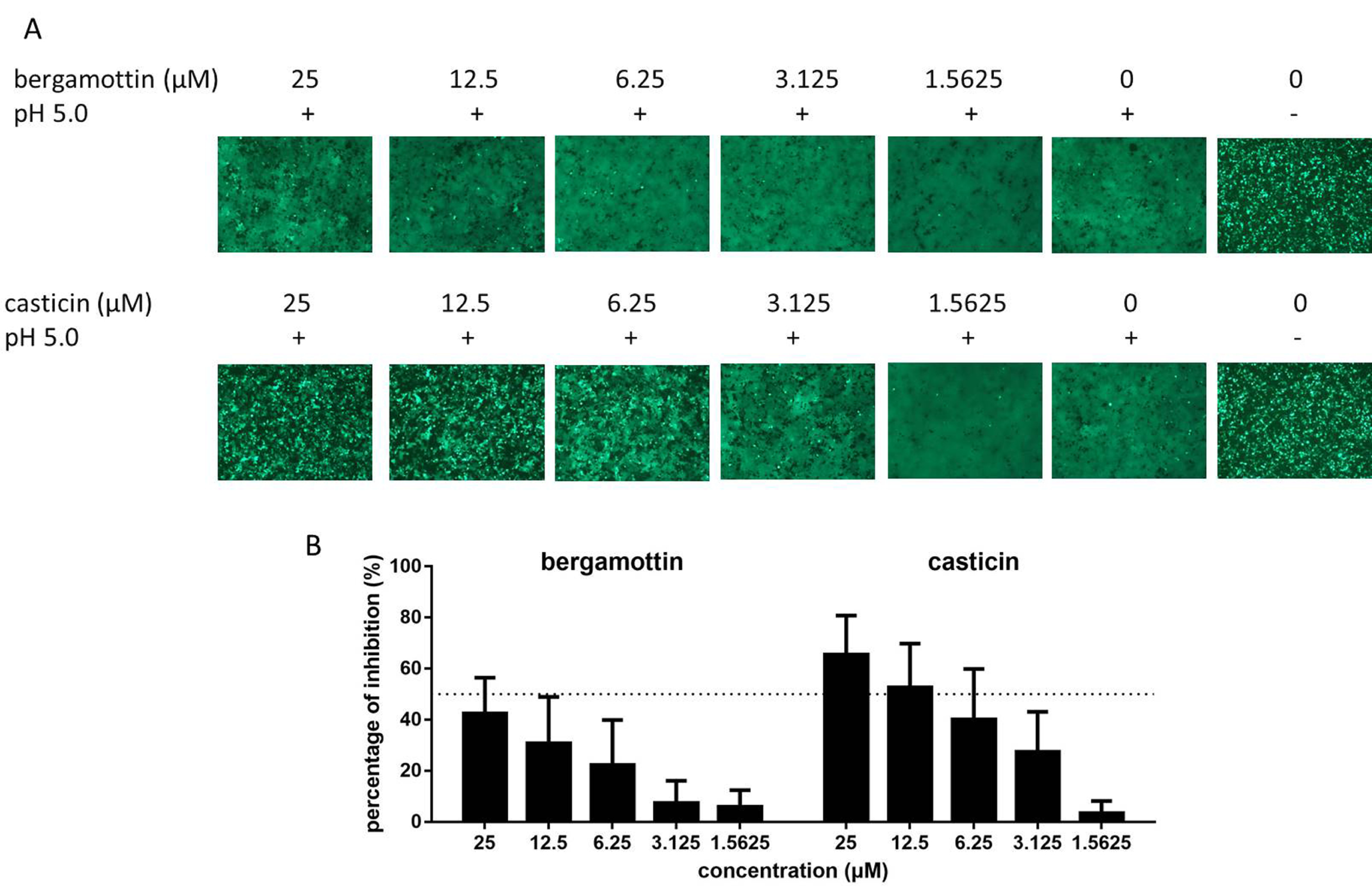
Bergamottin and casticin inhibited LASV GPC-mediated membrane fusion. (A) Qualitatively evaluate the inhibitory activities of bergamottin and casticin against membrane fusion. Syncytium formation was visualized by using fluorescent microcopy. Images are representative fields from 4 or 5 independent experiments. (B) A dual-luciferase assay was used to quantitatively evaluate the inhibitory activities of bergamottin and casticin against membrane fusion. Data are presented as means ± SDs for 3-4 independent experiments.

To further quantitatively evaluate the inhibitory activities, fusion efficacy was determined using the dual-luciferase assay (9, 11-13). As shown in Fig. 2B, casticin exhibited a dose-dependent inhibition of GPC-mediated membrane fusion. Furthermore, the inhibition effect shown in the membrane fusion assay was similar to that obtained from the infection assay (Fig. 1E). Notably, casticin could inhibit LASV GPC-mediated membrane fusion by ∼65% with 25 µM, while the inhibition by bergamottin was <50% at the same concentration. We increased the concentration of bergamottin to 100 µM, and the large-scale syncytium was still present, suggesting that bergamottin blocks LASV entry via a different mechanism from casticin.

### Bergamottin blocks LASV endocytic trafficking

To explore the mechanism underlying bergamottin inhibition, we further investigated the effects of the compounds on virus binding and endocytic trafficking. Firstly, binding efficacy was evaluated in the absence and presence of the compounds in A549 cells, and the IIH6 antibody (a blocking antibody targeting the glycan epitope on α-DG) was used as the positive control (14, 15). As shown in Fig. 3A, IIH6 significantly blocked LASVrv bound to the cell surface, while neither compound showed inhibitory effects, suggesting that both bergamottin and casticin did not interfere with the binding of LASV GP1 to the cell surface receptor.

**Fig. 3.**
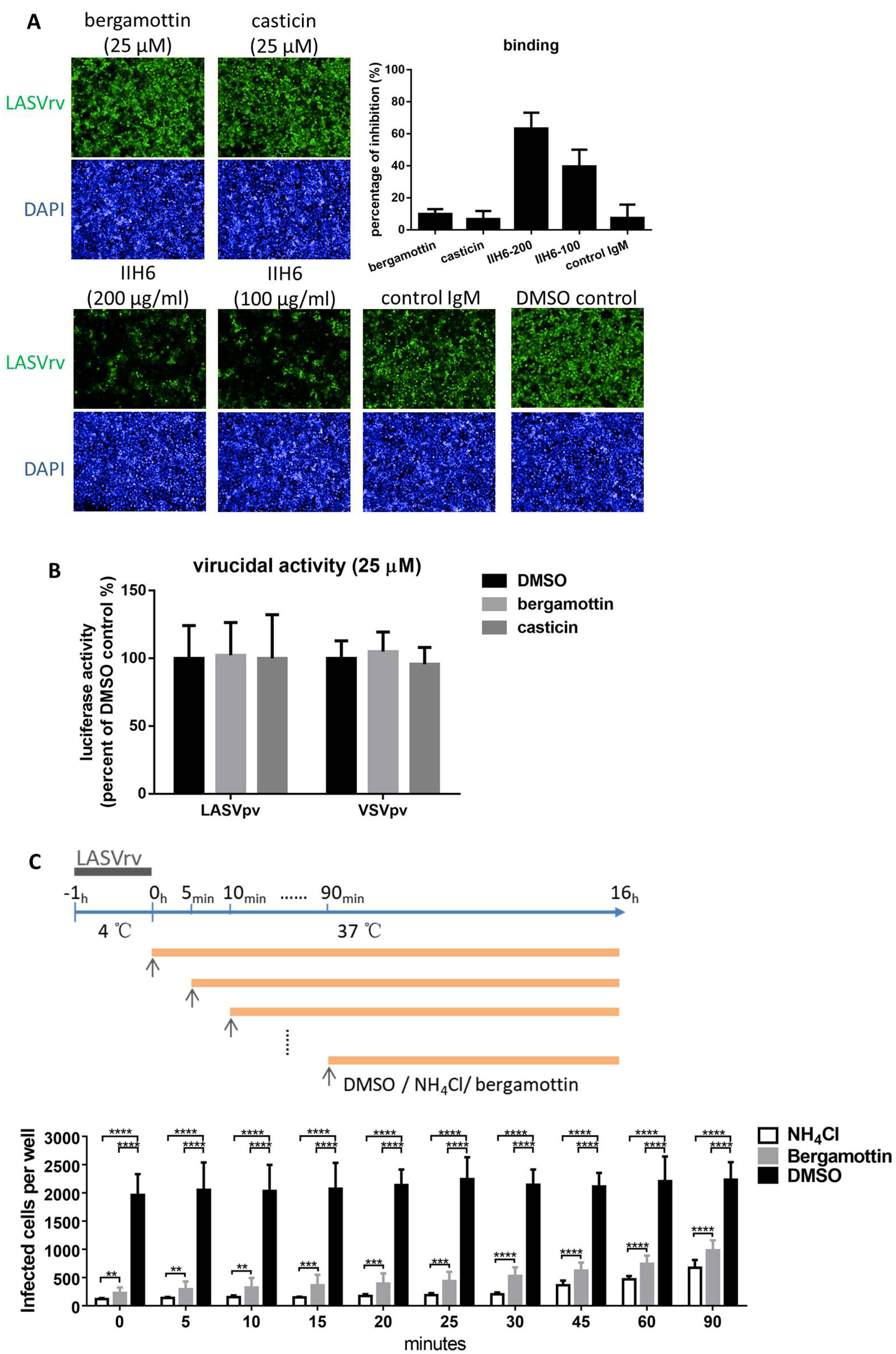
Bergamottin blocked LASVpv endocytic trafficking. (A) Effects of bergamottin and casticin on LASVpv binding. Quantitative analysis of the percentage of inhibition was based on the numbers of positive cells. Inhibition = (GFP_DMSO_/DAPI_DMSO_ - GFP_tested_/DAPI_tested_) / (GFP_DMSO_/DAPI_DMSO_). Error bars represent the SDs from three independent experiments. (B) Neither bergamottin nor casticin had virucidal effects. LASVpv or VSVpv at an MOI of 2 were incubated with DMSO or compounds (25 µM for 1 h) and then diluted 200-fold and added to cells. (C) Time-of-addition assay. The experimental timeline is shown in the upper panel. A549 cells were infected with LASVrv (MOI, 0.01) at 4°C for 1 h and then washed with PBS; the temperature was then increased to 37°C in the presence of bergamottin (25 µM) or NH_4_Cl (4.65 mM) for the indicated times. Data are presented as the means ± SDs for 5-8 independent experiments.

Next, we tested whether both compounds could bind to LASV GPC and exert a virucidal effect. To this end, LASVpv (2 × 10^5^ PFU) was mixed with either compound at 25 μM for 1 h; the mixture was then diluted 200-fold to the non-inhibitory concentration and added to the cells for 1 h (16). Luciferase activities were determined 23 h later. As shown in Fig. 3B, the luciferase activities observed in the compound treated groups were similar to those in the DMSO control group, suggesting that neither compound had a virucidal effect on LASVpv.

As bergamottin did not exert an inhibitory effect on the virucidal, binding, or fusion step, we hypothesized that it might inhibit the endocytic trafficking of the pseudotype of LASV. To address this, we quantitatively compared the inhibition of endocytic efficiency of bergamottin with the acidification antagonist NH_4_Cl. As shown in Fig. 3C, a 1-hour incubation with LASVrv (MOI of 0.01) at 4 °C led to approximately 2,000 cells being infected in a 96-well plate, while treatment with NH_4_Cl significantly decreased the number of infected cells to < 200, especially within the first 30 min. Similar to the inhibitory manner of NH_4_Cl, bergamottin suppressed the early stage of LASVrv infection. Although the inhibitory effect of bergamottin was relatively mild compared with that of NH_4_Cl, the most effective inhibition was found within the first 25-30 min. These results suggested that bergamottin inhibited LASV entry by acting on the endocytic trafficking of the virus. Notably, we have tried to select the adaptive mutants by serially passaging LASVrv in the presence of bergamottin, and no mutant was obtained after 12 rounds of passage, which was correlated with the results that bergamottin acted on the cellular endocytic pathway as opposed to targeting the virion.

### LASV GPC F446L mutation conferred resistance to casticin

Since most arenavirus fusion inhibitors serve as direct anti-viral agents by targeting the SSP-GP2 interface, we further investigated the viral targeting of casticin by selection for resistance mutations. As shown in Fig. 4A, after five rounds of passaging with 6.25 μM casticin (corresponding to an approximate IC_90_ value), LASVrv exerted a steady growth of resistance to casticin, while this was absent in DMSO control group. We sequenced the casticin resistant LASVrv from passage 6 (P6) to P10 without plaque purification. As shown in Fig. 4B, an adaptive mutant that changed from a phenylalanine (TTC) in the wild type to leucine (CTC) in the resistant virus, emerged from P7 and completed the substitution on P10. F446 is located at the transmembrane domain of GP2, which is highly conserved in mammarenaviruses (Fig. 4D). We introduced the F446L mutation into the GPC and generated a pseudotype of mutant LASV_F446Lpv_. As a result, LASV_F446Lpv_ conferred resistance to casticin. The inhibition efficiency against LASVpv WT infection reached ∼98% and ∼80% with casticin at 25 μM and 1.5625 μM, respectively, while only ∼15% inhibition against LASV_F446Lpv_ was observed when used with 25 μM casticin (Fig. 4C). Intriguingly, the F446 (corresponding to F438 in JUNV GPC) mutation has been reported to confer resistance to some structurally distinct compounds (17-20), suggesting that phenylalanine at this position might be accessible to different inhibitors, and the F446 mutation might contribute to the stabilization of GPC induced by the inhibitors.

**Fig. 4.**
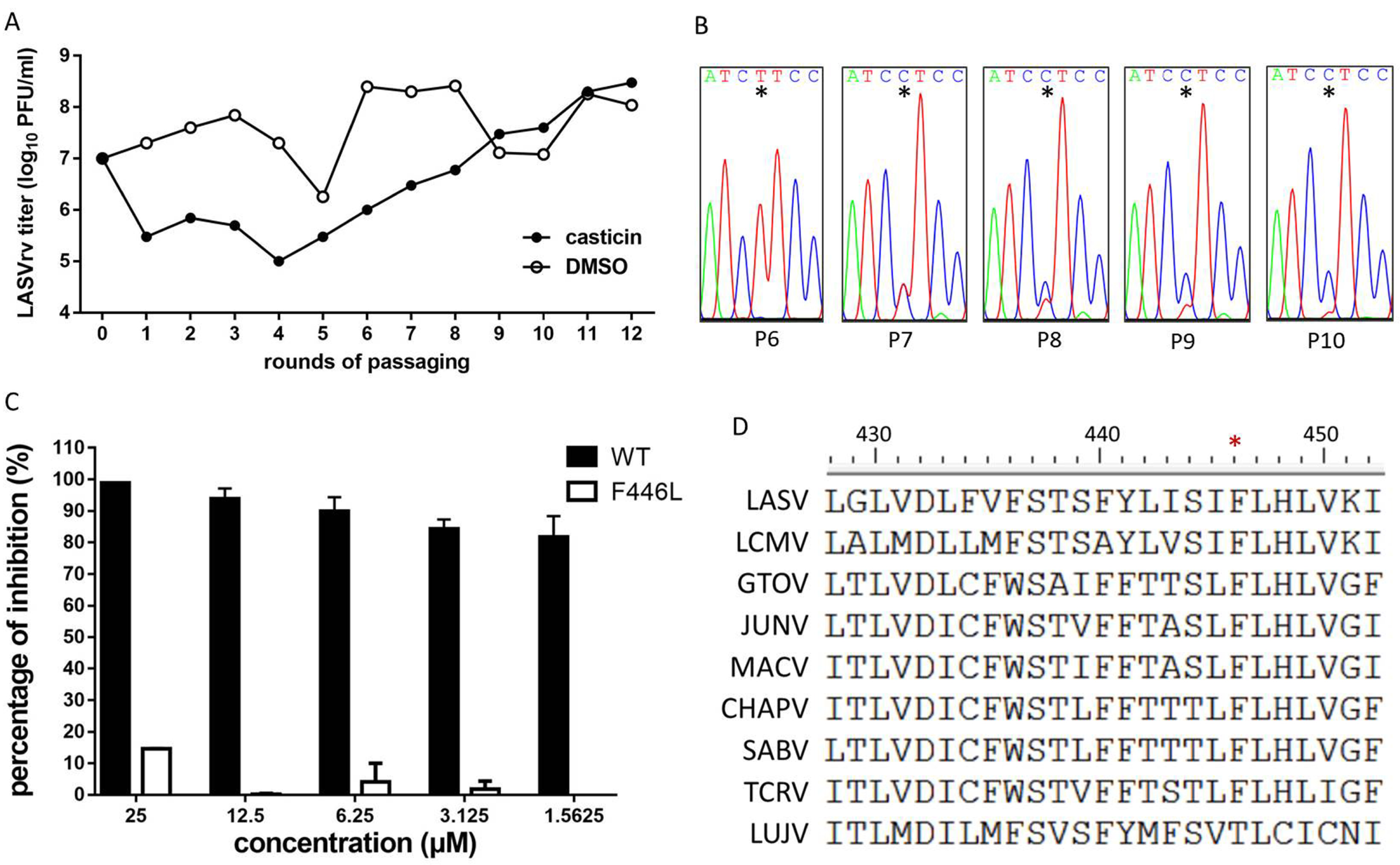
Selection of casticin-resistant LASVrv. (A) The adaptive mutant virus was selected by serially passaging LASVrv in the presence of 6.25 μM casticin. In a parallel experiment, LASVrv passaging in vehicle served as a control. (B) Sequencing chromatograms of P6 to P10 viruses. The asterisk highlights the emergence of the adaptive mutant. (C) Resistant activities of the WT and F446L LASVrv (MOI 0.01) to casticin. Data are presented as means ± SDs from 2-3 independent experiments. (D) Amino acid sequence alignment of the mammarenavirus transmembrane domain.

### Casticin extends the antiviral spectrum to other mammarenaviruses

As F446 is highly conserved in mammarenaviruses (Fig. 4D), we asked whether casticin could inhibit entry of other arenaviruses. To address this, pseudotypes of the pathogenic OW viruses (including the prototype lymphocytic choriomeningitis virus [LCMV], LUJV, and the most closely related Mopeia virus [MOPV]) and NW viruses (including JUNV, MACV, GTOV, SBAV, and CHAPV) were generated with the VSV backbone. As shown in Fig. 5, casticin could effectively inhibit both the NW and OW pseudotype virus infections in a dose-dependent manner. Moreover, we constructed the leucine mutant pseudotype of each virus with the corresponding F446 site mutation, and tested the sensitivities of these mutant viruses to casticin. Notably, all the mutant NW pseudotype viruses, in line with the LASV_F446Lpv_, conferred resistance to casticin, while the pathogenic OW viruses (LCMV and LUJV) bearing the leucine mutation were still sensitive to casticin. We supposed that casticin might inhibit LASV and NW mammarenaviruses entry by a similar mechanism. To this end, we tested whether casticin could inhibit the membrane fusion caused by NW GPC. As shown in Fig. 5B, casticin at 80 µM could inhibit NW GPC-mediated membrane fusion, while the corresponding F446L mutation conferred resistance to casticin inhibition by GPC-mediated membrane fusion.

**Fig. 5.**
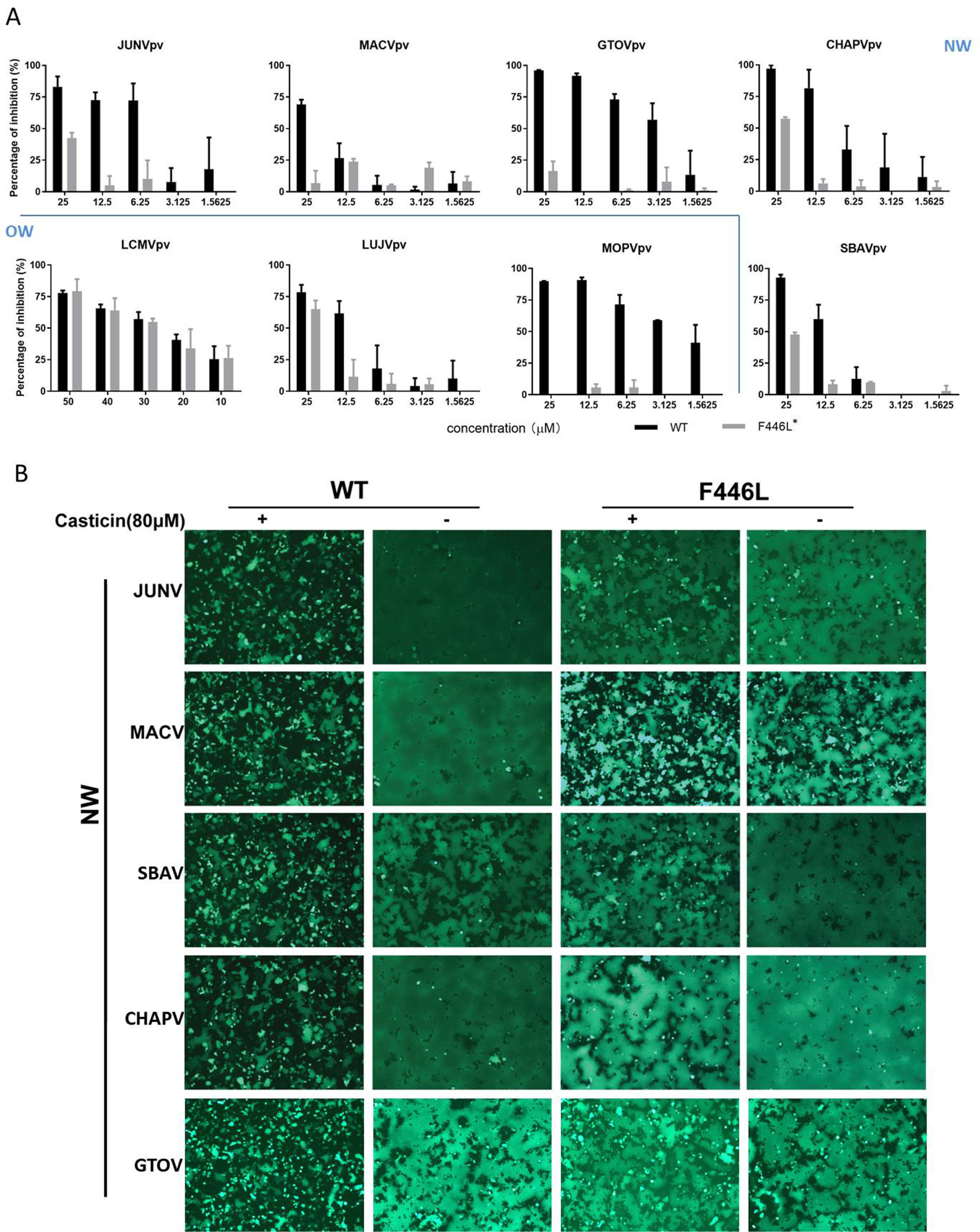
Casticin exerted the antiviral effects on NW mammarenaviruses. (A) Vero cells were incubated in the absence and presence of casticin. After 1 h, the pseudotypes of JUNV, MACV, GTOV, CHAPV, SBAV, LCMV, LUJV, MOPV, and the corresponding F446L mutant viruses were added at an MOI of 0.1. The supernatant was removed 1 h later, and the cell lysates were assessed for luciferase activity 23 h later. Data are presented as means ± SDs from 3 independent experiments. (B) Casticin inhibited NW GPC-mediated membrane fusion, while the conserved phenylalanine mutation conferred the resistance. 293T cells were transfected with pCAGGS-NW GPC; 24 h later, casticin (80 µM) was added for 1 h, followed by treatment with acidified (pH 5.0) DMEM for 15 min. Syncytium formation was visualized after 1 h. Images are representative fields from 3 independent experiments.

## Discussion

In this study, we screened a botanical drug library and identified two hit compounds, bergamottin and casticin, that prohibited the entry step of pseudotype and recombinant LASV infection. Bergamottin is a natural furanocoumarin found principally in grapefruit, which has been shown to inhibit cytochrome P450 (CYP450) enzymes and thus causes variations in the metabolism of many medications (21, 22). Herein, we showed that bergamottin inhibits LASVpv entry via blocking of viral endocytic trafficking, which seems to be irrespective of the native mechanism possessed by bergamottin. NW pathogenic arenaviruses use TfR1 as the primary receptor and consistently engage in clathrin-mediated endocytosis, in which the gradually lower pH leads to the maturation of the endosome (23). In contrast to NW arenaviruses, entry by the OW is initiated by binding with the primary receptor α-DG (24), bypassing the early endocytic pathway, and they are transported with the multivesicular bodies to the late endosome (25, 26). Notably, the OW specie LUJV utilizes the unusual NRP2 as the primary receptor that is needed to switch the receptor to CD63 to complete the productive entry (27, 28). Similarly, LASV undergoes receptor switching and engages LAMP1 in the late endosome for efficient fusion (14, 29). As both the NW and OW mammarenaviruses use receptor-mediated endocytosis to enter into cells, bergamottin is supposed to inhibit these viruses infections by blocking the endocytic trafficking. Notably, although cytochrome P450 enzymes are essential for the metabolism of many medications, the only FDA-approved drug for treatment of Lassa fever, ribavirin, is not a substrate of CYP450 enzymes (30). As ribavirin inhibits LASV by targeting the replication step, and bergamottin acts on the entry step, using a combination of both drugs might offer benefits over each as a single monotherapy. Similarly, the combined usage of bergamottin with other entry inhibitors, such as lacidipine, ST-161 and ST-193, etc., might provide synergistic antiviral effects.

We also identified casticin as a hit compound via HTS for its strong inhibition against LASV entry. Casticin is a constituent of *Vitex agnus-castus* (*agnus castus*) seeds; natural products containing casticin have been extensively used to treat premenstrual syndrome (PMS) with few adverse effects (31, 32). Intravenous injection of 50 mg/kg in rats has been shown to lead to a maximum plasma concentration (*Cmax*) of ∼34 µM (33), which is far higher than the IC_50_ reported here. Notably, casticin has also been shown to inhibit CYP3A4 enzyme (31, 32), which is the most abundant CYP enzyme in the liver and intestines. Herein, we demonstrated that casticin inhibits LASVpv entry by acting as a direct anti-viral agent (DAA). Casticin blocked the LASV GPC-mediated membrane fusion by targeting the SSP-GP2 interface, and the viral target was the conserved phenylalanine located in the transmembrane domain of GP2. Furthermore, we found that the phenylalanine to leucine mutation at the corresponding site caused the NW pathogenic mammarenaviruses to confer resistance to casticin. On the other hand, F446L is a high rate mutation that has been found to confer resistance to distinct entry inhibitors (17-19). Whether the phenylalanine at this site was accessible to the distinct compounds and thus participated in the compound-induced stabilization of the pre-fusion conformation of GPC, or the mutation at this site increases the stability of the glycoprotein needs further investigation.

Above all, the study identified bergamottin and casticin as LASV entry inhibitors, highlighting two hit compounds derived from natural products that worked with different mechanisms. This study also highlighted the role of F446 in the SSP-GP2 interface in the drug resistance in mammarenaviruses, and identified the broad-spectrum antiviral effects of casticin.

## MATERIALS AND METHODS

### Cells and viruses

BHK-21, HEK 293T, Vero, HeLa, and A549 cells were cultured in Dulbecco’s modified Eagle’s medium (DMEM; HyClone, Logan, UT, USA) supplemented with 10% fetal bovine serum (Gibco, Grand Island, NY, USA). The pseudotype VSV bearing the GPC of LASV (Josiah strain, GenBank accession number HQ688673.1), LCMV (Armstrong strain, AY847350.1), LUJV (NC_012776.1), MOPV (AY772170.1), GTOV (NC_005077.1), JUNV (XJ13 strain, NC_005081.1), MACV (Carvallo strain, NC_005078.1), SBAV (U41071.1), and CHAPV (NC_010562.1) were generated as previously reported (34, 35). 293T cells transfected with pCAGGSGPC were infected with pseudotype VSV (described below) in which the G gene was replaced with the luciferase gene at an MOI of 0.1 for 2 h. The culture supernatants were harvested 24 h later, centrifuged to remove cell debris, and stored at 80°C. The recombinant VSV expressing the GPC of LASV was generated as described previously (9, 35, 36). The plasmid used for constructing recombinant virus was pVSVΔG-eGFP (where eGFP is enhanced green fluorescent protein; plasmid 31842; Addgene). GPC was cloned into the ΔG site, and the construct was designated pVSVΔG-eGFP-GPC. BHK-21 cells in 6-well plates were infected with a recombinant vaccinia virus (vTF7-3) encoding T7 RNA polymerase at an MOI of 5. Forty-five minutes later, cells were transfected with 11 µg of mixed plasmids with a 5:3:5:8:1 ratio of pVSVΔG-eGFP-GPC (pVSVΔG-Rluc for generating pseudotype VSV), pBS-N, pBS-P, pBS-G, and pBS-L. After 48 h, the supernatants were filtered to remove the vaccinia virus and inoculated into BHK-21 cells that had been transfected with pCAGGS-VSV G 24 h previously. The pseudotype and recombinant viruses bearing LASV GPC are designated LASVpv and LASVrv, respectively. The titer of pseudotype virus was measured by infecting BHK-21 cells previously transfected with pCAGGS-VSV G and determined by plaque assay 24 h post infection. The titer of recombinant virus was determined by plaque assay. The titers of LASVpv and LASVrv were 3×10^7^/ml and 1.6×10^7^/ml, respectively.

### Optimization of HTS assay conditions

Cell density and MOI were optimized at 1 ×10^4^ cells and 1 × 10^2^ PFU per 96-well plate, respectively, as previously described (9, 37). Methyl-beta-cyclodextrin (MβCD; 2 mM) and 0.5% DMSO were used as the positive and negative controls, respectively. The signal-to-basal (S/B) ratio, coefficient of variation (CV), and Z’ factor were determined as 14269, 1.64% and 0.84, respectively (9, 37).

### HTS assay of the botanical drug library

A library of 1,058 compounds from botanical drugs was purchased from Weikeqi Biotech (Sichuan, China). Compounds were stored as 10 mM stock solutions until use. HTS was carried out as shown in Fig. 1A. Vero cells were treated in duplicate with the compounds (50 µM); 1 h later, cells were infected with LASVrv (MOI, 0.1). After 23 h, cells were fixed with 4% paraformaldehyde and stained with DAPI (4’, 6-diamidino-2-phenylindole) (Sigma-Aldrich, USA). Nine fields per well were imaged on an Operetta high-content imaging system (PerkinElmer), and the percentages of infected and DAPI-positive cells were calculated using the associated Harmony 3.5 software. Cell viability was evaluated by using 3-(4,5-dimethyl-2-thiazolyl)-2,5-diphenyl-2H-tetrazolium bromide (MTT) assay. To determine the IC_50_ values, LASVpv with MOI of 0.1 was used, and the timeline was conducted as above. The luciferase activity was measured using the Rluc assay system (Promega, Madison, WI, USA), and the IC_50_ was calculated using Graphpad Prism 6.

### Membrane fusion assay

293T cells co-transfected with pCAGGS-GPC and pEGFP-N1 were treated with compounds or vehicle (DMSO) for 1 h, followed by incubation for 15 min with acidified (pH 5.0) medium. The cells were then placed in neutral medium, and syncytium formation was visualized 1 h later.

For quantification of the luciferase-based fusion assay, 293T cells in 24-well plates transfected with both pCAGGS-GPC (0.25 µg) and plasmids expressing T7 RNA polymerase (pCAGT7, 0.25 µg) were cocultured at a ratio of 3:1 with targeted cells transfected with pT7EMCVLuc (2 µg per well for 6-well plates) and 0.1 µg pRL-CMV (plasmids used in this assay were kindly provided by Yoshiharu Matsuura, Osaka University, Osaka, Japan). After 12 h of incubation, the compound treatment and pH induction were conducted as described above. Cell fusion activity was quantitatively determined after 24 h by measuring firefly luciferase activity expressed by pT7EMCVLuc and was standardized with Rluc activity expressed by pRL-CMV by using the Dual-Glo luciferase assay (Promega) (27, 29, 30).

### Binding assay

A549 cells were pretreated with 25 µM bergamottin or casticin at 37°C for 1 h. As a control, A549 cells were pretreated with IIH6 (sc-53987; Santa Cruz) or a control IgM at 4 °C for 2 h in parallel (14, 15). The cells were then transferred onto ice, and LASVrv (MOI, 1) was added for 1 h. After being washed with cold phosphate-buffered saline (PBS) three times, the bound viral particles were imaged using an Operetta high-content imaging system (Perkin Elmer).

### Virucidal assay

To study the virucidal effects of the compounds, approximately 6.4 ×10^4^ PFU of LASVpv or VSVpv was incubated with compounds (25 µM) or vehicle at 37 °C for 1 h; the mixture was diluted 200-fold to a noninhibitory concentration (MOI, 0.01) to infect Vero cells in a 96-cell plate. Luciferase activity was determined 24 h later as described above.

### Time-of-addition assay

The experimental timeline is depicted in Fig. 3C. At time -1 h, A549 cells were infected with LASVrv (MOI, 0.01) at 4 °C for 1 h and washed with PBS; the temperature was then increased to 37 °C to synchronize the infections. Bergamottin (25 µM) was incubated with the cells for different time courses as shown in Fig. 3C.

### Selection of adaptive mutants

Drug-resistant viruses were generated by passaging LASVrv on Vero cells in the presence of 6.25 μM casticin. LASVrv was also passaged in the presence of 0.5% DMSO in parallel as a control. RNA from the resistant viruses was extracted using TRIzol (TaKaRa) and reverse transcribed using the PrimeScript RT reagent kit (TaKaRa). The GPC segment was amplified and sequenced as previously described (9). Mutant sites were introduced to LASVpv as previously described, and casticin sensitivities were determined by Rluc activities.

## ACKNOWLEDGEMENTS

We thank the Center for Instrumental Analysis and Metrology, Core Facility and Technical Support, and Center for Animal Experiment, Wuhan Institute of Virology, for providing technical assistance.

This work was supported by the National Key Research and Development Program of China (2018YFA0507204), the National Natural Sciences Foundation of China (31670165), Wuhan National Biosafety Laboratory, Chinese Academy of Sciences Advanced Customer Cultivation Project (2019ACCP-MS03), the Open Research Fund Program of the State Key Laboratory of Virology of China (2018IOV001).

